# Functional tradeoffs promote morphological disparity in the bat nasal cavity

**DOI:** 10.64898/2026.10.07.757456

**Authors:** Vaibhav Chhaya, Timothy D. Smith, Abigail A. Curtis, Thomas P. Eiting, Sharlene E. Santana

## Abstract

Multifunctional anatomical structures must accommodate potentially competing performance demands, leading to functional tradeoffs. The mammal rostrum (snout) is involved in multiple functions, including feeding, respiratory conditioning, olfaction, and–in most bats–echolocation. Therefore, it serves as an ideal system to investigate whether and how functional tradeoffs promote or constrain morphological diversification. While the external morphology of the rostrum has been linked with mechanical demands of feeding in bats, the factors and processes influencing its internal morphology, including the turbinals in the nasal cavity, remain comparatively unexplored. Here we investigate how internal rostral morphology reflects specialization and tradeoffs among respiratory air conditioning, echolocation, and olfaction by quantifying turbinal morphological diversity in a phylogenetically and ecologically diverse sample of bat species. We find that echolocation mode predominantly influences both the ethmoturbinal complex (ETc) and maxilloturbinal (MT) systems, with nasal-emitting bats having reduced and less complex turbinals compared to oral and non-laryngeally echolocating bat species. MT complexity and surface area are not associated with bioclimatic variables representing thermal variability and aridity. However, the ETc exhibits higher complexity in plantivorous bats than faunivorous bats, suggesting higher olfactory reliance. While MT morphological disparity was predominantly explained by echolocation mode, ETc morphological disparity was associated with both echolocation mode and diet, suggesting potentially opposing demands favoring turbinal reduction for sound transmission and elaboration for olfaction. Among nasal emitters, which must accommodate both demands within the nasal cavity, ETc complexity exhibited higher relative disparity compared to MT complexity. These patterns suggest that competing functional demands may promote or maintain morphological diversity, possibly through ecological specialization and/or many-to-one mapping between morphology and performance.

## Introduction

Organisms often use the same anatomical structure to perform multiple fitness-relevant functions. For example, avian bills contribute to feeding, thermoregulation, and vocal performance, fish swim bladders regulate buoyancy and can improve hearing ability, and insect antennae mediate various forms of chemosensation and mechanosensation (Friedman et al., 2019; Krishnan & Sane, 2015; Ladich & Schulz-Mirbach, 2016). Evolutionary theory predicts that performance cannot be simultaneously maximized across multiple functions because physical principles, developmental pathways, and phylogenetic constraints impose tradeoffs among competing functions (Gould & Lewontin, 1979; Irschick & Higham, 2015). Such tradeoffs are thought to shape phenotypic diversity by favoring trait combinations that enhance performance in specific ecological contexts (Higham et al., 2021; Koehl, 1996). Under the traditional view, functional trade-offs are thought to constrain phenotypic diversification by limiting the range of phenotypes capable of performing multiple functions adequately (Corn et al., 2021; Muñoz, 2019; Polly, 2020; Stayton et al., 2018; Wainwright, 2007). However, this view assumes that the adaptive landscape integrating performance across functions is relatively simple, with a single adaptive peak in phenotypic space (Arnold, 2003; Stayton et al., 2018). When the adaptive landscape instead contains multiple peaks, either because multiple regions of phenotypic space confer similar performance for a given function (i.e., many-to-one mapping) or when lineages exhibit ecological specialization, functional tradeoffs may promote or retain phenotypic disparity instead (Alfaro et al., 2005; Shoval et al., 2012; Stayton et al., 2018; Wainwright et al., 2005). Multifunctional anatomical structures thus offer an excellent opportunity to test whether functional tradeoffs constrain or promote phenotypic diversification because their morphology reflects the performance demands of multiple, potentially competing functions.

The bat rostrum (snout) combines multifunctionality with exceptional morphological diversity, making it a powerful system for examining the effects of functional tradeoffs on morphological diversification. Bats (order Chiroptera) are among the most ecologically diverse mammalian groups, varying widely in habitat, diet, and sensory ecology (Santana et al., 2025). This diversity places varied performance demands on the rostrum, which is involved in food acquisition and mastication, respiratory conditioning, olfaction, and echolocation, among several other fitness-relevant functions (Adams & Pedersen, 2013; Page & ter Hofstede, 2021; Santana et al., 2025). Across Chiroptera, much of the external cranial shape variation is explained by changes in rostrum morphology, particularly rostral length and orientation about the braincase (Arbour et al., 2019). Early shifts in rostral orientation strongly match the evolution of echolocation modes (loss, oral, and nasal), and subsequent changes in rostral length are associated with dietary evolution (Arbour et al., 2019). The link between external rostrum shape and diet is especially pronounced and has been mechanistically studied in phyllostomid bats, where species that consume tough foods have evolved shorter rostra that can generate greater bite forces due to higher mechanical advantage (Dumont et al., 2011; Santana et al., 2012; Santana et al., 2010). However, little is known about how variation in the morphology of the bat rostrum, particularly its internal anatomy, relates to the multiple sensory and physiological functions it performs, or how these potentially competing functional demands have influenced its morphological evolution.

In most tetrapods, a major source of rostral internal morphological diversity lies in the turbinals, intricate bony and cartilaginous structures that occupy the nasal passages (Martinez et al., 2024; Negus, 1958). Turbinals are largely covered by either respiratory or olfactory mucosa, which are responsible for air conditioning (regulating the heat and moisture of respired air) and olfaction, respectively (Negus, 1958; Smith et al., 2015). In mammals, the respiratory turbinals, predominantly the maxilloturbinals (MT), are situated anteriorly in the nasal cavity and directly in the path of respiratory airflow, where they facilitate heat and water exchange through a counter-current system (Jackson & Schmidt-Nielsen, 1964; Owerkowicz et al., 2015; Schmidt-Nielsen et al., 1970) (Figure 1A). Because the magnitude of this exchange depends on surface area, species inhabiting extreme environments (e.g., xeric habitats or cold climates) are expected to show the most architecturally complex respiratory turbinals with the greatest relative surface area (Baldo et al., 2026; Cheon et al., 2023; Mason et al., 2020; Van Valkenburgh et al., 2011), however there is limited evidence supporting this idea (Green et al., 2012; Martinez et al., 2023; Van Valkenburgh et al., 2004). The olfactory turbinals, comprising the ethmoturbinals, frontoturbinals, and interturbinals, are located more posteriorly within the nasal cavity, often partially contained within a cul-de-sac known as the olfactory recess (Negus, 1958; Smith et al., 2015) (Figure 1A, S2). Greater surface area and complexity of these turbinals have been associated with increased olfactory ability, as inferred from diet, habitat, and other ecological proxies (Green et al., 2012; Martinez et al., 2018, 2020; Yohe et al., 2022). However, the specific aspects of olfactory performance (e.g., odor sensitivity versus discrimination) that benefit from this morphology remain unclear (Yohe et al., 2022). The two functional classes of turbinals discussed above often show differences in their relative surface area that are correlated with ecological requirements in different mammal groups. In smaller-bodied species and species with shortened snouts, spatial tradeoffs may arise between respiratory and olfactory turbinals because the rostrocaudal space available constrains the total surface area that can be accommodated within the nasal cavity (Martinez et al., 2018, 2020; van Valkenburgh et al., 2014; Zwicker et al., 2018).

**Figure 1.**
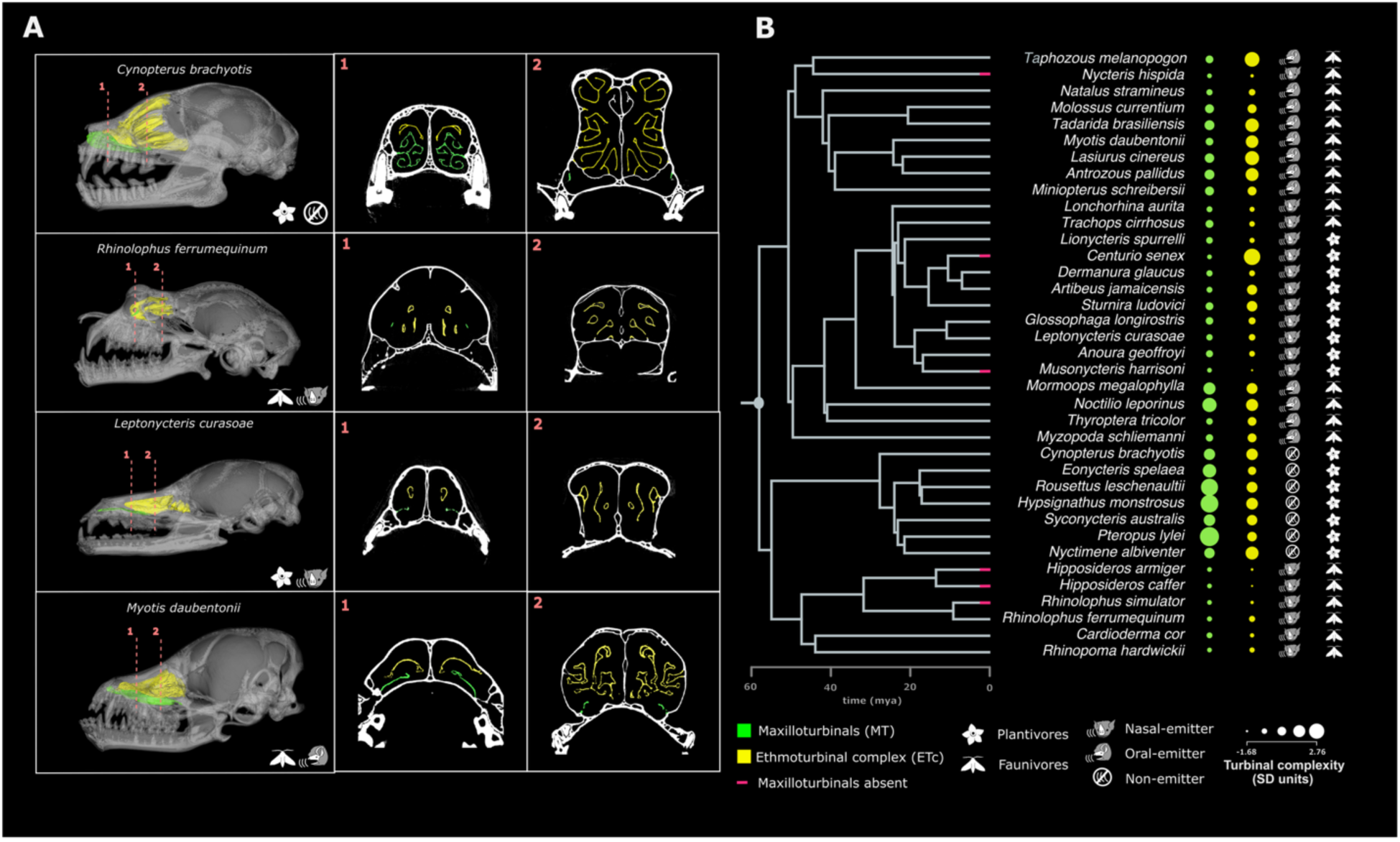
**(A)** Three-dimensional reconstructions and representative coronal CT sections of the maxilloturbinals (MT; green) and ethmoturbinal complex (ETc; yellow) in four bat species. **(B)** MT and ETc complexity, diet, and echolocation mode mapped onto the maximum-clade-credibility tree for the species included in this study (Upham et al., 2019). Circle size represents turbinal complexity expressed in standard deviation units.

Most bats emit echolocation calls through their mouth (‘oral-emitters’), but at least four lineages have convergently evolved the use of their nasal passages for echolocation signal emission (‘nasal-emitters’) (Pedersen, 1993, 1998; Pedersen & Müller, 2013) (Figure 1). Anatomical descriptions report reduction or loss of turbinal elements in some nasal-emitting bats, with unique strand-like modifications of the ethmoturbinals in rhinolophid bats hypothesized to contribute to echolocation (Curtis et al., 2020; Curtis & Simmons, 2017; King et al., 2026). Although the link between sound transmission and turbinal morphology in mammals remains unexplored in a comparative context, surgical reduction of hypertrophied turbinals in humans has been shown to lead to significant changes in nasalance and perceived vocal quality, indicating that turbinal morphology may affect the properties of emitted sounds (Valsamidis et al., 2022). Additionally, bat species that do not use laryngeal echolocation (‘non-emitters’, family Pteropodidae) possess more structurally complex and numerous turbinals than laryngeally echolocating bats, including both oral- and nasal-emitting species (King et al., 2026). Together, these findings suggest that echolocation mode may alter the functional demands placed on the nasal passages of bats (Dzal & Gillam, 2024; Pedersen, 2000), which must therefore balance respiratory air conditioning and olfaction with the additional acoustic requirements potentially imposed by nasal echolocation.

We quantify respiratory and olfactory turbinal morphological diversity in a phylogenetically and ecologically diverse sample of bat species to investigate how potentially competing sensory and physiological functions may have shaped the evolution of internal rostral morphology. Using micro-computed tomography data and phylogenetic comparative methods, we test two overarching hypotheses. First, we hypothesize that turbinal morphology reflects the sensory and physiological demands associated with respiratory air conditioning, olfaction, and echolocation across ecological contexts. Under this hypothesis, we predict that (1) respiratory turbinals, specifically the maxilloturbinals (MT), exhibit greater complexity and relative surface area in species inhabiting more thermally variable and xeric environments than in those inhabiting more thermally stable and humid environments, reflecting enhanced heat and water conservation (Baldo et al., 2026; Cheon et al., 2023; Green et al., 2012; Martinez et al., 2023; Mason et al., 2020; Van Valkenburgh et al., 2011) ; (2) olfactory turbinals, grouped as the ethmoturbinal complex (ETc), exhibit greater complexity and surface area in plantivorous species (frugivores and nectarivores) than in faunivorous species (insectivores and carnivores), consistent with greater reliance on olfaction to locate fruits and flowers (Brokaw et al., 2021; Denzinger & Schnitzler, 2013; Gonzalez-Terrazas et al., 2016; Korine & Kalko, 2005; Luft et al., 2003; Raghuram et al., 2009; Smith et al., 2011; Thies et al., 1998; von Helversen et al., 2000); and (3) both MT and ETc complexity and surface area are reduced in nasal-emitting species, potentially improving sound transmission by altering the acoustic filtering properties of the nasal passages (Curtis et al., 2020; Dzal & Gillam, 2024; King et al., 2026). Second, we hypothesize that tradeoffs among respiratory air conditioning, olfaction, and echolocation influence morphological diversification. If functional tradeoffs constrain the range of morphologies capable of adequately performing multiple functions, we predict lower relative disparity in turbinal morphology in nasal-emitting bats than in oral-emitting and non-laryngeally echolocating bats. This prediction would reflect potential performance tradeoffs between nasophonation and respiratory conditioning in the MT, and nasophonation and olfaction in the ETc.

## Methods

### Micro-CT scanning and segmenting of bony turbinals

To quantify three-dimensional turbinal morphology, we analyzed micro-computed tomography (µCT) scans of one skull specimen each from 37 bat species, spanning 16 of the 21 currently recognized chiropteran families (Morales et al., 2026) (Figure 1B). Our dataset combined previously published µCT scans (Arbour et al., 2019, 2021; Grossnickle et al., 2024) with new scans acquired for this study (Supplementary Data). Specimens were loaned from the Burke Museum of Natural History and Culture (Seattle, WA) and the American Museum of Natural History (New York, NY), and scanned using a Bruker SkyScan 1172 µCT scanner (Bruker, Billerica, USA; Supplementary Data). CT slices were reconstructed from raw projection data using NRecon software (v1.7.4.6, Bruker, Billerica, USA).

From these CT image stacks, we segmented the full cranium of each specimen by manually thresholding bone from air and soft tissue across all slices, then used regional thresholding to aid in segmentation of the turbinal skeleton in Mimics software (v23.0; Materialise, Leuven, Belgium). To account for size-related effects on turbinal morphology, we measured cranial length from the segmented cranium as the distance between the anterior-most midpoint of the internasal suture (rhinion) and the dorsal-most point of the foramen magnum (opisthion; Figure S1A). We also calculated rostrum volume (approximated as palatal length × palatal width × rostrum depth) as a more anatomically localized alternative to cranial length to account for size effects, given that the rostrum houses the turbinals and exhibits the greatest morphological disparity in the bat skull (Arbour et al., 2019). Palatal length was measured between the anterior and posterior ends of the palatal midline; palatal width was measured between the lingualmost points of the left and right last molars in palatal view; and rostrum depth was measured as the perpendicular distance from the dorsalmost point of one half of the cribriform plate to the palatal midline (Figure S1B, C).

Then, we segmented the bony turbinal structures in each specimen. Segmented structures were assigned to two turbinal systems, the maxilloturbinals (MT) and the ethmoturbinal complex (ETc), which serve as functional proxies for respiratory air conditioning and olfaction, respectively (King et al., 2026; Martinez et al., 2020; Negus, 1958; Smith et al., 2015; Van Valkenburgh et al., 2004, 2011) (Figure 1A). Maxilloturbinals were identified based on their rostroventral position within the nasal cavity proper and their articulation with the maxillary bone. The ethmoturbinal complex included the ethmoturbinals and interturbinals, which articulate with the ethmoid bone; the frontoturbinals, which articulate with the frontal bone; and the horizontal lamina, a bony shelf descending from the lateral nasal wall that often anchors to the ethmoturbinals and interturbinals (Ito et al., 2021; King et al., 2026; Maier, 1993; Smith et al., 2015) (Figure S2). We excluded other bony laminae, such as the semicircular and transverse laminae, from the ETc because they are absent in some bat species and contribute minimally to olfactory epithelial surface area in most bats (King et al., 2026) . However, we included the horizontal lamina because it frequently supports olfactory epithelium (e.g., see Figure 5 in King et al., 2026). Although the ethmoturbinals, particularly ET I, may exhibit some respiratory epithelial coverage rostrally, most of the ETc surface area supports olfactory epithelium (King et al., 2026; Smith et al., 2012, 2024). We therefore use ETc surface area as a proxy for olfactory epithelial coverage. For both MT and ETc segmentations, if turbinal elements were fused with adjacent laminae, we segmented only the portions projecting into the nasal cavity lumen.

### Quantifying turbinal surface area and complexity

We used two metrics to quantify turbinal morphology: turbinal surface area and turbinal complexity. To measure surface area, we generated 3D surface meshes of the segmented maxilloturbinals (MT) and ethmoturbinal complex (ETc) and extracted surface areas for each region in Mimics software. We exported MT and ETc surfaces as STL files from Mimics and imported them into MorphoDig (ver. 1.6.7; Lebrun, 2018) to quantify turbinal complexity. Using MorphoDig, we used the ‘*Fit convex hull’* function to generate a convex hull enclosing each turbinal structure and subsequently used the ‘*Measure surface area’* function to calculate the surface area of the corresponding convex hull.

We calculated turbinal complexity for the MT and ETc of each species as:

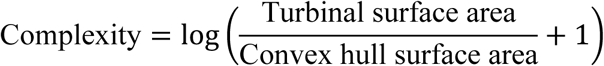

This complexity metric represents the amount of turbinal surface area relative to the rostro-caudal space that the turbinals occupy. A species lacking bony turbinals would show a turbinal complexity of 0, while a species with geometrically complex turbinals would have a complexity closer to 1.5 (the maximum observed value of complexity).

### Dietary, sensory, ecological, and behavioral data

To evaluate olfactory, echolocation, and air conditioning performance as predictors of turbinal morphology, we used diet, echolocation mode, and bioclimatic variables, respectively, as the ecological and behavioral proxies for these functional demands.

We adopted two broad dietary categories reflecting the relative reliance on olfaction for detecting and localizing food sources: (1) species that primarily consume fruits and nectar (“plantivores”) and (2) species that primarily consume insects or other animals (“faunivores”), both based on prior dietary assessments (Arbour et al., 2019; Rajemison & Goodman, 2007; Wilman et al., 2014). Omnivorous species (*Anoura geoffroyi, Antrozous pallidus, Glossophaga longirostris, Trachops cirrhosus*) were classified based on whether plants or animals constituted the largest proportion of their diet. Although both dietary groups likely integrate multiple sensory modalities during foraging, plantivores are known to rely on olfactory cues to initially detect and localize fruits or flowers, whereas faunivores primarily depend on passive hearing or echolocation to locate animal prey (Brokaw et al., 2021; Denzinger & Schnitzler, 2013; Gonzalez-Terrazas et al., 2016; Korine & Kalko, 2005; Luft et al., 2003; Raghuram et al., 2009; Thies et al., 1998; von Helversen et al., 2000).

We categorized species into three echolocation modes: species that do not echolocate laryngeally ("non-emitters"), species that emit echolocation calls through the nasal passages ("nasal-emitters"), and species that emit calls through the mouth ("oral-emitters"), following previous assessments (Pedersen, 1993; Pedersen & Müller, 2013). Although this classification is widely used, we recognize that vocal behaviors in bats are complex and context-dependent. For example, members of Phyllostomidae, a predominantly nasal-emitting family, have been observed flying with their mouths open, raising the possibility that oral emission may occur in some contexts (Gessinger et al., 2021). Conversely, three genera within Vespertilionidae, a primarily oral-emitting family, have been reported to emit calls nasally, although none of these species were included in our sample (Jakobsen et al., 2018; Seibert et al., 2015). Nevertheless, these exceptions do not alter the predominant emission mode of each species, and the three-category classification provides a useful framework for testing broad evolutionary patterns across bats (Arbour et al., 2019).

As proxies of respiratory air-conditioning demands, we extracted bioclimatic variables from the WorldClim database using specimen locality coordinates and the R packages *terra* and *geodata* (Fick & Hijmans, 2017; Hijmans, 2026; Hijmans et al., 2026). We focused on two bioclimatic variables with potential relevance to respiratory heat and water exchange: BIO7 (temperature annual range) and BIO17 (precipitation of the driest quarter). BIO7 was used as a proxy for thermal variability, under the assumption that species inhabiting environments with greater seasonal temperature fluctuations may experience stronger respiratory heat exchange demands. BIO17 was used as a proxy for aridity because lower precipitation during the driest part of the year is generally associated with an increased risk of respiratory water loss. These variables provide broad indicators of climatic conditions that may influence the air conditioning demands placed on the maxilloturbinals. For most species, climatic values were extracted from the locality at which the scanned specimen was collected at a resolution of 10-arc minutes. One exception was *Hypsignathus monstrosus*, for which the specimen originated from a captive colony at the Bronx Zoo, New York. For this species, we extracted climatic values across its native IUCN distribution and used the maximum BIO7 and minimum BIO17 values observed within the range to approximate the upper bounds of thermal variability and aridity that the species may encounter in the wild (Tanshi, 2016).

### Phylogenetic analyses of turbinal morphology

To test for the effect of echolocation, diet and climate on turbinal morphology, we fitted phylogenetic generalized least squares (PGLS) models with a Pagel’s λ correlation structure, using the functions ‘gls’ (package *nlme*) and ‘corPagel’ (package *ape*) (Pagel, 1999; Paradis & Schliep, 2019; Pinheiro et al., 2026) in RStudio (ver 2026.09.0; R Core Team, 2026). For our phylogenetic analyses, we used a set of 1,000 phylogenetic trees pruned to our species sample from VertLife.org (Upham et al., 2019). We fitted separate models for MT complexity, MT surface area, ETc complexity, and ETc surface area as response variables. MT and ETc surface areas were transformed as log (surface area + 1) before analysis, with the constant added to accommodate zero values in species lacking bony turbinals (Supplementary Data). Echolocation mode was included as a predictor in all models. The MT models additionally included annual temperature range (BIO7) and precipitation of the driest quarter (BIO17), whereas the ETc models included diet. Log-transformed cranial length was included as a covariate in all models to control for size-related effects on turbinal morphology. Additionally, we fitted these same models with log-transformed rostrum volume instead of cranial length.

For models incorporating cranial length in which ETc complexity or log-transformed ETc surface area were the response variables, maximum-likelihood estimation of λ consistently failed to converge. Examination of likelihood profiles generated by fitting these models across a range of fixed λ values across all trees indicated that the likelihood function peaked at the lower boundary of the parameter space (λ = 0) (Figure S3). Consequently, λ was fixed at 0 for these analyses, representing no phylogenetic signal in the residuals and effectively reducing the models to generalized least squares (GLS) regressions with uncorrelated errors. For models incorporating cranial length in which MT complexity or MT surface area were the response variables, λ was successfully estimated by maximum likelihood and was used to account for phylogenetic non-independence among species. For models incorporating rostrum volume, likelihood profiles peaked at λ = 0 for all four response variables; consequently, λ was fixed at 0 for all these analyses (Figure S4).

For all models, we performed Type II ANCOVA tests (with log-transformed cranial length or rostrum volume as the covariate) to assess the significance of predictors using the ‘Anova’ function (package *car*), followed by post hoc pairwise comparisons for significant effects using Tukey contrasts implemented via the ‘glht’ function (package *multcomp*) (Fox & Weisberg, 2019; Hothorn et al., 2008). If MT morphology reflects demands associated with respiratory air conditioning, then MT complexity and surface area should increase with annual temperature range (BIO7) and decrease with precipitation during the driest quarter (BIO17). If ETc morphology reflects olfactory demands associated with diet, then plantivorous species should exhibit greater ETc complexity and surface area than faunivorous species. Finally, if nasal emission constrains internal nasal morphology, then nasal-emitting species should exhibit lower MT and ETc complexity and surface area than oral-emitters and non-emitters.

Following the associations identified by the PGLS analyses, we fitted a series of continuous-trait evolutionary models to determine whether differences in MT and ETc complexity among echolocation modes and dietary groups were consistent with distinct selective regimes. We used the functions ‘mvBM’ and ‘mvOU’ (package *mvMORPH*) to fit three model classes: single-regime Brownian motion (BM1), single-regime Ornstein–Uhlenbeck (OU1), and multi-regime Ornstein– Uhlenbeck (OUM) (Clavel et al., 2015). If dietary and echolocation mode categories represent distinct selective regimes, then multi-regime OUM models should outperform the single-regime BM1 and OU1 models. The OUM model allows regime-specific phenotypic optima (θ) while constraining both the evolutionary rate (σ²) and the strength of selection (α) to be constant across regimes. For MT complexity, regimes were defined by the three echolocation mode categories (OU3), whereas for ETc complexity, we additionally fit a two-regime model with the two dietary categories as the selective regimes (OU2). For the multi-regime models, we generated 10 stochastic character maps for each topology across a random subsample of 100 trees from the initial posterior distribution, using the ‘make.simmap’ function in the package *phytools* (Revell, 2024). These mappings were based on an ‘equal rates’ model (ER), which provided the best fit for both echolocation mode and diet evolution compared to the ‘symmetric rates’ (SM) and ‘all-rates-different’ (ARD) models (Kruskal-Wallis, *p* < 0.001). Model fit was evaluated by comparing median small-sample-size-corrected Akaike Information Criterion (AICc) values and corresponding Akaike weights across all the stochastic character mappings. Models with ΔAICc < 2 were considered to have substantial support and were treated as the best-fitting models. We tested whether the median AICc values were significantly different from each other using a Kruskal-Wallis test, implemented by the ‘kruskal.test’ function (R Core Team, 2026). Additionally, we calculated and compared the proportion of stochastic character mappings for which each model was identified as the best-fitting model. Finally, to assess whether our model selection procedure had sufficient power to discriminate among candidate models given our sample size and regime distribution, we conducted parametric bootstrap analyses (Boettiger et al., 2012). We estimated median parameters for the best-fitting model across our subsample of trees and used these to simulate 500 replicate datasets using the ‘mvSIM’ function, with regimes mapped onto the maximum clade credibility tree from our posterior distribution. Candidate models were refit to each replicate using the selection procedure described above, and a Kruskal-Wallis test was used to assess whether AICc scores differed significantly among models. These tests detected significant differences among candidate models for both MT and ETc (Kruskal-Wallis, *p* < 0.001).

Finally, we quantified morphological disparity in MT and ETc complexity as the sample variance, calculated using the ‘var’ function (R Core Team, 2026). If competing functional demands constrain morphological diversification, then the turbinal system experiencing stronger competing demands should exhibit lower relative disparity. Alternatively, greater relative disparity would indicate that these demands can be accommodated through multiple morphological solutions. Variance was calculated across all species and separately within echolocation subgroups. For each subgroup, relative morphological disparity was calculated as the within-subgroup variance divided by the total variance across all species for the corresponding turbinal system. Thus, the relative disparity for subgroup g is given by 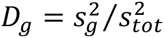.

## Results

### Echolocation mode predicts maxilloturbinal morphology independent of climate

PGLS models revealed that maxilloturbinal (MT) complexity is strongly and significantly predicted by echolocation mode when cranial length is used as a size covariate (ANCOVA, *p* < 0.001) (Table 1, Figure 2A). Specifically, nasal-emitting bat species exhibit significantly lower MT complexity than both oral-emitting and non-emitting species (Tukey’s HSD, adjusted *p* < 0.001 for nasal-oral and nasal-none comparisons). Several nasal-emitting taxa from Rhinolophidae, Hipposideridae, Phyllostomidae, and Nycteridae lack bony maxilloturbinals, indicating convergent loss of this structure across multiple independent lineages (Figure 1B); MT loss was not observed in any oral- or non-emitting species. Oral-emitting species display intermediate MT complexity, whereas non-emitting species (all belonging to the family Pteropodidae) exhibit the highest MT complexity (Figure 1B, 2A). However, these two groups were not significantly different from each other after accounting for other predictors in the PGLS model (Tukey’s HSD, adjusted *p* = 0.788 for oral-none comparison). In contrast, climate variables BIO7 (temperature annual range) and BIO17 (precipitation of the driest quarter), our proxies for climatic extremeness, had no significant effect on MT complexity (ANCOVA, median *p* = 0.631 and 0.619 respectively) (Figure 2A). These patterns remained the same when MT surface area was analyzed as the response variable instead of complexity (Table 1, Figure 2A).

**Figure 2.**
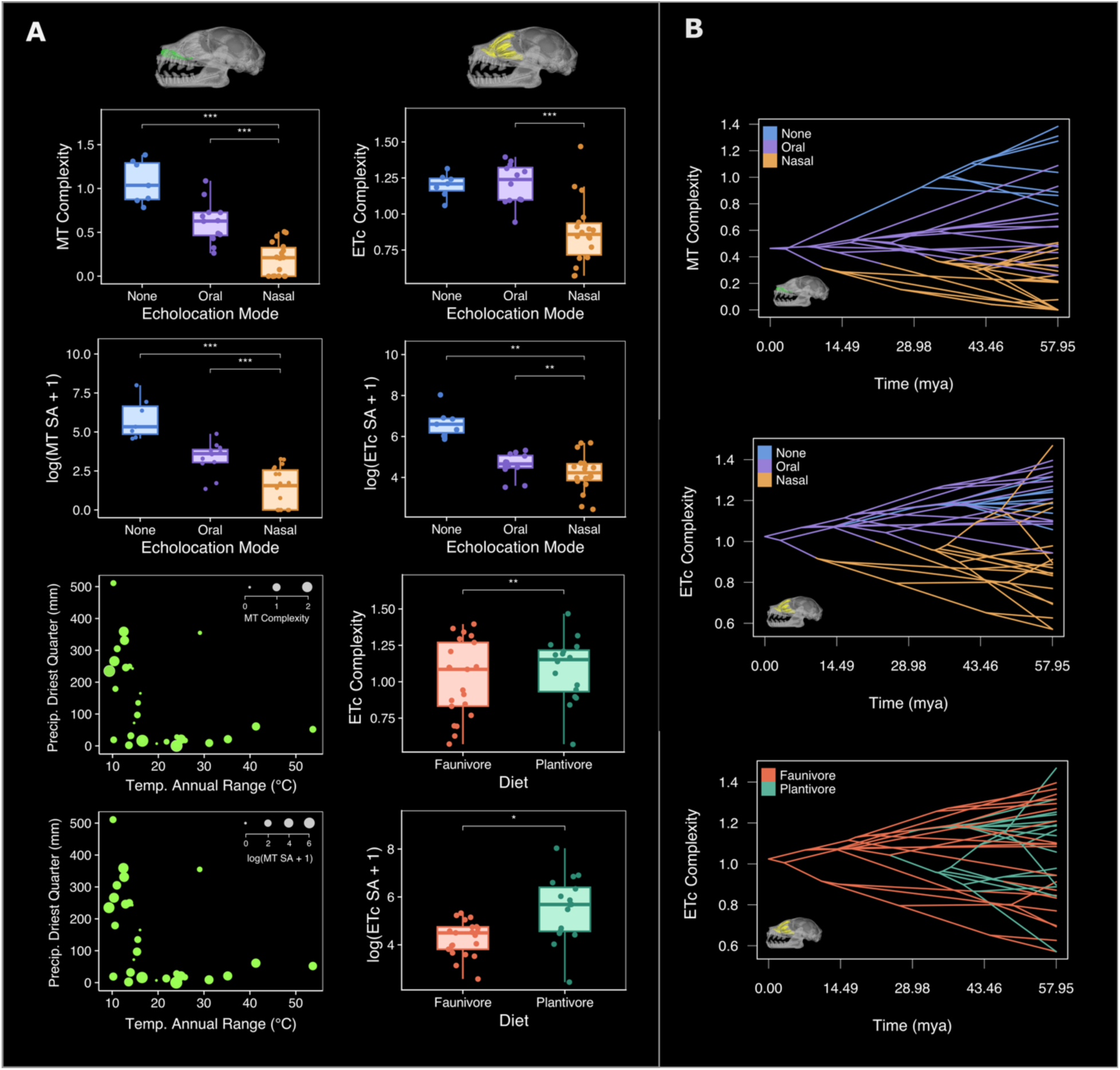
**(A)** Boxplots showing maxilloturbinal (MT) and ethmoturbinal (ETc) complexity and surface area across echolocation modes and dietary categories. Asterisks indicate significance from ANCOVA tests of phylogenetic generalized least-squares (PGLS) models: *p < 0.05, **p < 0.01, and ***p < 0.001. Echolocation mode significantly predicted MT and ETc complexity and surface area, whereas diet significantly predicted ETc complexity and surface area. Neither annual temperature range nor precipitation during the driest quarter significantly predicted MT complexity or surface area. **(B)** Phenograms (Revell, 2024) illustrating the diversification of MT and ETc complexity. In each phenogram, the phylogeny is projected into trait space, with species positioned according to their observed complexity values and branches representing inferred changes in complexity through evolutionary time. Branch colors represent stochastic character maps of echolocation mode for MT and ETc complexity and of diet for ETc complexity.

**Table 1.** Phylogenetic generalized least squares (PGLS) ANCOVA results for maxilloturbinal (MT) complexity and surface area using log-transformed cranial length as the size covariate. Models included precipitation of the driest quarter (BIO17), annual temperature range (BIO7), and echolocation mode as predictors. Pairwise comparisons among echolocation modes were conducted using Tukey’s HSD post hoc test. Results are summarized as medians across phylogenetic trees; percentage support indicates the proportion of trees for which each effect or comparison was significant (p < 0.05).

| **PGLS model: MT complexity ~ BIO 17 + BIO 7 + Echolocation mode + log (Cranial length)** |  |  |  |
| --- | --- | --- | --- |
| **ANCOVA summary** |  |  |  |
| **Predictor** | **Median $\chi^2$** | **Median p-value** | **% support across trees** |
| BIO 17 (Precipitation of the driest quarter) | 0.25 | 0.619 | 0 |
| BIO 7 (Annual temperature range) | 0.23 | 0.631 | 0 |
| log (Cranial length) | 7.88 | 0.005 | 100 |
| Echolocation mode | 62.59 | <0.001 | 100 |
| **Tukey's HSD summary (Predictor: Echolocation mode)** |  |  |  |
| **Comparison** | **Median contrast estimate** | **Median adjusted p-value** | **% support across trees** |
| Nasal - None | -0.683 | <0.001 | 100 |
| Nasal - Oral | -0.566 | <0.001 | 100 |
| Oral - None | -0.117 | 0.788 | 0 |
| **PGLS model: log (MT surface area+1) ~ BIO 17 + BIO 7 + Echolocation mode + log (Cranial length)** |  |  |  |
| **ANCOVA summary** |  |  |  |
| **Predictor** | **Median $\chi^2$** | **Median p-value** | **% support across trees** |
| BIO 17 (Precipitation of the driest quarter) | 0.119 | 0.730 | 0 |
| BIO 7 (Annual temperature range) | 0.001 | 0.971 | 0 |
| log (cranial length) | 8.142 | 0.004 | 100 |
| Echolocation mode | 45.036 | <0.001 | 100 |
| **Tukey's HSD summary (Predictor: Echolocation mode)** |  |  |  |
| **Comparison** | **Median contrast estimate** | **Median adjusted p-value** | **% support across trees** |
| Nasal - None | -3.442 | <0.001 | 100 |
| Nasal - Oral | -2.677 | <0.001 | 100 |
| Oral - None | -0.765 | 0.725 | 0 |

Both MT complexity and surface area showed a significant positive relationship with rostrum volume (ANCOVAs, *p* < 0.001) (Table S1). When log-transformed rostrum volume was used as the size covariate, MT complexity and surface area remained significantly associated with echolocation mode (ANCOVA, *p* < 0.001 for both), and climate variables remained as non-significant predictors (ANCOVA, *p* > 0.05 for BIO7 and BIO17) (Table S1). Nasal-emitting species exhibit significantly lower MT complexity and surface area for a given rostral volume than both oral-emitting and non-emitting species (Tukey’s HSD, adjusted *p* < 0.01 for all nasal-oral and nasal-none comparisons), with no significant differences between the latter two groups after accounting for rostral volume (Tukey’s HSD, adjusted *p* > 0.5 for oral-none comparisons) (Table S1).

Evolutionary model-fitting analyses supported the strong influence of echolocation mode on MT complexity, where a three-optimum OU (OUM) model corresponding to the three echolocation modes consistently exhibited the lowest AICc score than both the single-regime Brownian motion (BM1) and single-optimum OU (OU1) models across all stochastic character mappings (median ΔAICc relative to OUM = 14.00 and 19.93 respectively) (Table 3) (Figure 2B). This indicates that MT complexity has evolved toward distinct trait optima associated with the three echolocation modes, rather than a single optimum shared across all species.

### Echolocation mode predicts ethmoturbinal complex morphology more strongly than diet

Ethmoturbinal complex (ETc) complexity and surface area were also significantly predicted by echolocation mode after accounting for log-transformed cranial length (ANCOVA, p < 0.001 for both), with nasal-emitting species generally exhibiting significantly lower complexity and surface area values than both oral-emitting and non-emitting bats (Tukey’s HSD, adjusted *p*: nasal-oral< 0.01 for both complexity and surface area, nasal-none = 0.05 for complexity and 0.004 for surface area) (Table 2). Dietary category was also a significant predictor of ETc morphology, although its effect was weaker than that of echolocation. Plantivorous species overall exhibited greater ETc complexity than faunivorous species, whereas differences in ETc surface area were less pronounced (ANCOVA, p = 0.002 and 0.046 respectively) (Table 2).

**Table 2.** Phylogenetic generalized least squares (PGLS) ANCOVA results for ethmoturbinal complex (ETc) complexity and surface area using log-transformed cranial length as the size covariate. Models included diet and echolocation mode as predictors. Pairwise comparisons between dietary categories and among echolocation modes were conducted using Tukey’s post hoc test.

| **PGLS model: ETc complexity ~ Diet + Echolocation mode + log (Cranial length)** |  |  |  |  |
| --- | --- | --- | --- | --- |
| **ANCOVA summary** |  |  |  |  |
| **Predictor** | **$\chi^2$** | **Median p-value** |  |  |
| Diet | 9.406 | 0.002 |  |  |
| Echolocation mode | 43.996 | <0.001 |  |  |
| log (Cranial length) | 0.348 | 0.555 |  |  |
| **Tukey's HSD summary (Predictor: Diet)** |  |  |  |  |
| **Comparison** | **Contrast estimate** | **Lower 95% CI** | **Upper 95% CI** | **Adjusted p-value** |
| Plantivore - Faunivore | 0.243 | 0.088 | 0.399 | 0.002 |
| **Tukey's HSD summary (Predictor: Echolocation mode)** |  |  |  |  |
| **Comparison** | **Contrast estimate** | **Lower 95% CI** | **Upper 95% CI** | **Adjusted p-value** |
| Oral - None | 0.190 | -0.146 | 0.525 | 0.366 |
| Nasal - None | -0.246 | -0.493 | 0.001 | 0.050 |
| Nasal - Oral | -0.436 | -0.615 | -0.257 | <0.001 |
| **PGLS model: log (ETc surface area+1) ~ Diet + Echolocation mode + log (Cranial length)** |  |  |  |  |
| **ANCOVA summary** |  |  |  |  |
| **Predictor** | **$\chi^2$** | **Median p-value** |  |  |
| Diet | 3.976 | 0.046 |  |  |
| Echolocation mode | 24.825 | <0.001 |  |  |
| log (Cranial length) | 4.090 | 0.043 |  |  |
| **Tukey's HSD summary (Predictor: Diet)** |  |  |  |  |
| **Comparison** | **Contrast estimate** | **Lower 95% CI** | **Upper 95% CI** | **Adjusted p-value** |
| Plantivore - Faunivore | 0.679 | 0.012 | 1.347 | 0.046 |
| **Tukey's HSD summary (Predictor: Echolocation mode)** |  |  |  |  |
| **Comparison** | **Contrast estimate** | **Lower 95% CI** | **Upper 95% CI** | **Adjusted p-value** |
| Oral - None | -0.429 | -1.869 | 1.010 | 0.751 |
| Nasal - None | -1.460 | -2.519 | -0.401 | 0.004 |
| Nasal - Oral | -1.030 | -1.799 | -0.261 | 0.005 |

ETc surface area was significantly associated with rostrum volume (*p* < 0.001), but ETc complexity was not (*p* = 0.395) (Table S2). When log-transformed rostrum volume was used as a covariate, echolocation mode remained a significant predictor of both ETc complexity and surface area (ANCOVA, *p* < 0.001 and *p* = 0.003 respectively) (Table S2). Pairwise comparisons for ETc complexity indicated that oral-emitting species had significantly greater complexity for a given rostral volume than both nasal-emitting and non-emitting species (Tukey contrasts, adjusted *p* < 0.001 and *p* = 0.046 respectively), whereas nasal-emitting and non-emitting species did not differ significantly (adjusted *p* = 0.365) (Table S2). For ETc surface area, nasal-emitting species exhibit significantly lower values than both oral-emitting and non-emitting species (adjusted *p* = 0.035 and 0.031 respectively). Plantivorous species have significantly greater ETc complexity than faunivorous species (ANCOVA, *p* = 0.013), but the dietary effect on ETc surface area was not significant after accounting for rostral volume (*p* = 0.272) (Table S2).

Consistent with the patterns demonstrated by our regressions, the three-optimum OU model based on echolocation mode provided a substantially better fit than both BM1 and OU1, indicating that ETc evolution is better described by distinct adaptive optima associated with echolocation mode than by a single evolutionary optimum shared across all species (Table 3). In contrast, the two-optimum OU model based on dietary regimes (OU2) was the poorest fit of all candidate models.

**Table 3.** Evolutionary model comparisons for maxilloturbinal (MT) and ethmoturbinal complex (ETc) complexity. Models included single-rate Brownian motion (BM1), single-optimum Ornstein-Uhlenbeck (OU1), and Ornstein-Uhlenbeck models with distinct optima for dietary categories (OU2) or echolocation modes (OU3). Model fits were evaluated using the small-sample corrected Akaike information criterion (AICc) and Akaike weights, summarized across stochastic character maps. Percentage support indicates the proportion of maps for which each model had the lowest AICc.

| **Evolutionary model fits: MT complexity** |  |  |  |  |
| --- | --- | --- | --- | --- |
| **Model** | **Median AICc** | **ΔAICc** | **Median Akaike weight** | **% support across simmaps** |
| BM1 | 12.59 | 14.00 | 0.001 | 0 |
| OU1 | 18.57 | 19.93 | <0.001 | 0 |
| **OU3 (echolocation)** | **-1.45** | **0.00** | **0.999** | **100** |
| **Evolutionary model fits: ETc complexity** |  |  |  |  |
| **Model** | **Median AICc** | **ΔAICc** | **Median Akaike weight** | **% support across simmaps** |
| BM1 | -1.62 | 13.40 | 0.001 | 0 |
| OU1 | -0.10 | 15.00 | 0.001 | 0 |
| OU2 (diet) | 1.95 | 17.00 | <0.001 | 0 |
| **OU3 (echolocation)** | **-15.00** | **0.00** | **0.998** | **100** |

### Morphological disparity differs between maxilloturbinals and the ethmoturbinal complex across echolocation modes

Across all the species sampled, MT complexity exhibited greater variance (s²ₜₒₜₐₗ = 0.161) than ETc complexity (s²ₜₒₜₐₗ = 0.060). Within nasal-emitting bats, however, ETc complexity showed considerably higher relative morphological disparity (Dₙₐₛₐₗ = 0.865) than MT complexity (Dₙₐₛₐₗ = 0.223) (Figure 2B). The difference between the two turbinal systems was considerably smaller within the other echolocation groups; oral emitters exhibited nearly identical relative disparity in MT and ETc complexity (Dₒᵣₐₗ = 0.358 and 0.337, respectively), while non-echolocating bats exhibited moderately greater relative disparity in MT complexity than in ETc complexity (Dₙₒₙₑ = 0.369 and 0.118, respectively) (Figure 2B).

## Discussion

Our comparative study strongly suggests that diversification of the internal rostral morphology of bats is shaped by an interplay of multiple functional demands. Echolocation mode is the main predictor of both respiratory (maxilloturbinal) and olfactory (ethmoturbinal complex) morphology, with nasal-emitting species exhibiting convergent reduction in turbinal complexity and relative surface area compared to oral-emitting and non-echolocating bats. Additionally, olfactory turbinal morphology is associated with diet (a proxy for olfactory reliance), with plantivorous species possessing greater complexity and relative surface area than faunivores. Respiratory turbinal morphology was not associated with climatic variables that reflect thermal variability and aridity, despite the importance of thermoregulation and water conservation for small mammals such as bats. Finally, the ethmoturbinal complex exhibited greater relative morphological disparity than the maxilloturbinals, consistent with the possibility that interactions among multiple functional demands can promote, rather than constrain, morphological diversification.

Nasal emission has evolved independently multiple times in bats, primarily in the superfamily Rhinolophoidea and the families Nycteridae and Phyllostomidae (Pedersen, 1993; Pedersen & Müller, 2013). All these lineages (except for frugivorous Phyllostomids) exhibit a striking simplification of their turbinals, possessing simple plate-like projections rather than the elaborate scrolled structures seen in other bats. This pattern is particularly marked in the respiratory maxilloturbinals (MT), which in many species are reduced to a single dorsally projecting structure or are absent, consistent with previous histological and diceCT descriptions (Curtis & Simmons, 2017; Ito et al., 2021; King et al., 2026). Evolutionary model comparisons further supported the association between echolocation mode and turbinal simplification: models allowing distinct trait optima among echolocation modes best explained both MT and ETc complexity. Together with the repeated reduction of turbinals in lineages that independently evolved nasal-emission, these results are consistent with convergence toward echolocation-associated morphological optima. Because thermoviscous losses are generally greater in narrower passages with higher wall surface area relative to airway volume, such convergent simplification of turbinals may enhance echolocation capabilities in nasal-emitting species by decreasing acoustic transmission losses and/or by modifying the acoustic filtering properties of the nasal passages (Kinsler et al., 2000). In many nasal-emitting species, the simplified MT merges with the semicircular lamina and frequently partitions greatly enlarged lateral recesses from the main nasal cavity (King et al., 2026). Although the function of these paranasal recesses remains unknown, they may contribute to acoustic filtering by acting as side-branch resonators and altering frequency bandwidths relevant to echolocation requirements (Ma et al., 2016; Suthers et al., 1988). Future work should use computational acoustic models to directly test how turbinal reduction and the associated evolution of lateral recesses influence sound transmission and frequency filtering, and whether these modifications improve echolocation performance.

Oral-emitting bats, although possessing more complex turbinals than nasal-emitting species, still exhibit lower respiratory (MT) turbinal complexity than non-echolocating bats. Although these differences were not statistically significant after accounting for phylogenetic relationships, they nevertheless raise the possibility that echolocation also exerts a functional demand on nasal morphology in oral-emitting taxa. Whether nasal- and oral-emitting bats produce biosonar signals exclusively through the nose and mouth, respectively, remains questionable, particularly in light of evidence suggesting that the mode of sound emission may be behaviorally flexible in some families (Gessinger et al., 2021; Seibert et al., 2015). If oral-emitting species can facultatively engage the nasal passages as a part of their vocal tract filter using velopharyngeal coupling (as documented in many other mammals), they may experience functional demands like those acting on nasal-emitting taxa, albeit to a lesser extent (Beeck et al., 2022; Fant, 1971; Koda et al., 2018; Reby et al., 2018; Sanvito et al., 2007). This possibility raises intriguing questions about the behavioral mechanisms that control sound production in bats, an area that remains remarkably understudied (Chen et al., 2025).

Besides the strong effect of echolocation mode, diet explained additional variation in olfactory turbinal (ETc) morphology in our sample. Plantivorous bats (primarily frugivorous and nectarivorous species) possessed more elaborate turbinals than faunivorous species (insectivorous and carnivorous species), potentially reflecting greater olfactory ability. Prior comparative studies of phyllostomid bats also showed that plant-visiting species possess greater olfactory turbinal surface area than faunivorous species (Hutcheon et al., 2002; Yohe et al., 2022), and our findings suggest that this pattern extends across a broader phylogenetic scale. However, despite the significant association with diet, evolutionary model comparisons did not support a model with separate ETc complexity optima for plantivores and faunivores, instead favoring optima associated with echolocation mode. This suggests that diet contributes to variation in ETc morphology within a more deeply rooted evolutionary pattern predominantly driven by echolocation mode, like that observed for the evolution of cranial shape across bats (Arbour et al., 2019). Overall, patterns of ETc morphological evolution are consistent with the possibility that functional demands associated with nasophonation and olfaction compete in laryngeally echolocating bats.

Although we were unable to statistically test differences among finer dietary categories because of sample size, we observed that the olfactory turbinals of nasal-emitting nectarivores were strikingly reduced compared with those of nasal-emitting frugivores, which had some of the highest ETc complexity values in our sample (e.g., *Centurio senex*, *Sturnira ludovici*; Figure S5). This contrast between nasal-emitting frugivores and nectarivores is unexpected because both rely on plant-produced odorants to locate their food (Korine & Kalko, 2005; Thies et al., 1998; von Helversen et al., 2000) and exhibit similar olfactory airflow patterns within the nasal cavity (but different flow rates in the ETc region) (Eiting et al., 2015, 2023). Additionally, nectarivores have elongated rostra that could still potentially accommodate substantial turbinal surface and complexity as nasal-emitting frugivores do in their foreshortened rostra, making the reduced turbinal complexity of nectarivores particularly difficult to explain simply by spatial constraints.

The trends observed among plantivorous species suggest that ETc morphology alone may provide an incomplete measure of olfactory reliance. Anatomical studies in bats and other mammalian taxa show that the olfactory epithelium extends onto other nasal structures, such as the nasal septum, so ETc surface area alone does not capture the total area available for odor detection (Smith et al., 2004, 2011, 2012). At the cellular and molecular levels, variation in olfactory epithelial thickness, olfactory sensory neuron density, olfactory receptor diversity, and the relative abundance of neurons expressing particular receptors could potentially compensate for reduced turbinal complexity (Eiting et al., 2023). Additionally, airflow patterns shaped by gross nasal cavity morphology and modulated by sniffing behavior may influence odorant delivery to the sensory epithelium (Eiting et al., 2014, 2015). Integrating turbinal morphology with epithelial anatomy, computational airflow modeling, behavioral experiments, and genetic approaches will therefore be essential for understanding how these features collectively shape olfactory function in bats.

An interplay between potentially competing functions was not detected for the respiratory turbinals (MT), as echolocation mode was the predominant predictor of MT complexity and surface area. Climate variables representing temperature variability and aridity were not associated with MT morphology as predicted, adding to evidence that MT morphology in mammals may not primarily reflect adaptation to environmental conditions or thermoregulatory demands (Green et al., 2012; Hillenius, 1992; Martinez et al., 2023; Schmid, 1976; Van Valkenburgh et al., 2004). Although the respiratory epithelium covering the MTs may contribute to respiratory air conditioning, bats possess numerous other mechanisms that may compensate for reduced thermoregulatory and osmoregulatory performance resulting from MT reduction. For example, in addition to the surface area occupied by respiratory epithelium in the nasal cavity, the efficiency of heat and water exchange depends on the architecture of the underlying vascular networks. Denser vascular networks and prevalence of venous sinuses may permit effective exchange even with the limited surface area offered by simple or absent bony turbinals, as may be the case in the nasal-emitting family Hipposideridae (Smith et al., 2022). Compensatory counter-current exchange may also occur in the trachea, as in birds, which have relatively simpler turbinals than mammals but longer tracheae and greater tracheal dead space (Hinds & Calder, 1971; Owerkowicz et al., 2015). Bats may also employ other physiological and behavioral strategies, ranging from selecting thermoneutral roosting sites to torpor, hibernation, estivation, migration, and exchanging heat through their highly-vascularised wings (Kluger & Heath, 1970; Rubalcaba et al., 2022). This redundancy in thermoregulation and water conservation mechanisms may weaken the relationship between maxilloturbinal morphology and climatic conditions, allowing the functional demands of nasophonation to exert a strong selective influence on the MT.

Collectively, our tests of functional associations with turbinal morphology revealed an interesting contrast between MT and ETc evolution. While the evolution of MT morphology was predominantly associated with echolocation mode, ETc morphology was associated with both echolocation mode and diet. ETc may therefore face potentially opposing selective pressures: reduction associated with nasophonation may facilitate echolocation while compromising olfactory performance. When we examined complexity among nasal emitters, ETc retained much of the variance observed across all species, whereas MT exhibited a much narrower distribution. Thus, rather than converging on an intermediate morphology representing the ‘best compromise’ between functions, nasal emitters occupied a broad range of ETc complexity values. A functional trade-off may therefore promote or maintain ETc morphological disparity rather than constrain it. This finding opposes the traditional view that multiple competing functional demands limit the range of phenotypes available (Corn et al., 2021; Muñoz, 2019; Polly, 2020; Stayton et al., 2018; Wainwright, 2007). One mechanism that could produce our observed pattern is ecological specialization toward alternative resolutions of the trade-off. Theory predicts that differences in the relative fitness contributions of competing functions can favor different ‘optimal’ compromises (Polly, 2020; Shoval et al., 2012). Because the relative importance of olfactory and acoustic performance appears to vary among nasal-emitting lineages, selection may not favor a single intermediate ET morphology (Gonzalez-Terrazas et al., 2016; Korine & Kalko, 2005; Leiser-Miller et al., 2020; Muchhala et al., 2024; Thies et al., 1998; von Helversen et al., 2000). Species that depend more strongly on olfaction (e.g., frugivorous phyllostomids) may retain elaborate ethmoturbinal complexes despite potential costs on echolocation performance, whereas those experiencing greater demands on echolocation performance and reduced use of olfaction while foraging (e.g., insectivorous rhinolophids) may exhibit ETc reduction. In contrast, MT morphology may face a narrower range of viable solutions under the predominant influence of nasophonation, resulting in stronger convergence toward reduced complexity. Another possible mechanism could involve many-to-one mapping and compensatory variation in other components of the olfactory and echolocation systems (as discussed above), allowing distinct ETc morphologies to achieve similar combined performance (Wainwright, 2007; Wainwright et al., 2005). Such cases of morpho-functional redundancy are prevalent, ranging from the feeding performance of centrarchid and labrid fish jaws to hindlimb morphology and jumping performance in *Anolis* lizards (Alfaro et al., 2005; Toro et al., 2004). Besides functional drivers, developmental differences between the two turbinal groups may also contribute to differences in their morphological diversification patterns. While the MT largely comprises a single turbinal element, the ETc contains several elements that can vary in their presence, relative size, and elaboration. Developmental studies of bat turbinals have documented differences in the growth trajectories of these individual turbinal elements, as well as changes in their configuration through splitting and fusion (Ito et al., 2021; Smith et al., 2024). Such modularity, although untested, could provide additional, intrinsic routes to diversification of the ETc and potentially contribute to its greater relative disparity among nasal emitters.

In conclusion, our findings suggest that morphological diversification in multifunctional structures is shaped by a complex interplay of functional demands. Anatomical structures subject to competing functional demands need not be constrained to a narrow region of phenotypic space (Corn et al., 2021; Muñoz, 2019; Polly, 2020; Stayton et al., 2018; Wainwright, 2007). Rather, ecological specialization and many-to-one mapping between morphology and performance may retain or even facilitate morphological diversification, depending on the topography of the adaptive landscape (Alfaro et al., 2005; Arnold, 2003; Polly, 2020; Shoval et al., 2012; Stayton et al., 2018; Wainwright et al., 2005). Our comparative study therefore challenges the traditional view that functional tradeoffs necessarily constrain morphological diversification and highlights the need for future work to explicitly quantify the performance of multifunctional structures across competing functions.

## Data and code availability

All data and code used in the analyses, along with associated specimen information, are provided in the supplementary data. CT scans and STL files used in this study will be deposited in MorphoSource (https://www.morphosource.org) upon publication.

## Acknowledgements and funding info

We thank Jeff Bradley at the Burke Museum of Natural History and Culture, Seattle, and Marisa Surovy at the American Museum of Natural History, New York, for their assistance with specimen loans. We thank Dr. Nancy Simmons for providing access to the CT scan of *Hypsignathus monstrosus*, and Chris Law, Dave Grossnickle, Annie Colgan, and Vivek Hari Sridhar for their helpful feedback and methodological suggestions. This study was funded by a National Science Foundation grants (award #2202271 to SES, #2202272 to TDS and #2336218 to TPE)

