## Supplementary Code for "Functional tradeoffs promote morphological disparity in the bat nasal cavity": Turbinals_analysis.R

```
# Libraries
library(ape)
library(car)
library(tidyverse)
library(phytools)
library(phangorn)
library(picante)
library(nlme)
library(mvMORPH)
library(multcomp)
library(emmeans)

# Replace spaces in species names with underscores
underscorer = function(sp_vector){
  separate = strsplit(sp_vector,' ')
  combined = lapply(separate,function(x) paste(x,collapse="_"))
  unlist(combined)
}

## reading tree object
batstrees <- read.nexus(file = "batstrees.nex")
batstrees_mcc <- maxCladeCred(batstrees)

## reading and prepping data

turbinal_data <- read.csv(file = "turbinal_data.csv")
turbinal_data$species <- underscorer(turbinal_data$species)
rownames(turbinal_data) <- turbinal_data$species
turbinal_data$diet <- as.factor(turbinal_data$diet)
turbinal_data$echolocation <- factor(turbinal_data$echolocation,c("None","Oral","Nasal"))
turbinal_data$diet_2cat <- as.character(turbinal_data$diet)
turbinal_data$diet_2cat[c(which(turbinal_data$diet == "Insectivore" | turbinal_data$diet =="Carnivore"))] = "Faunivore"
turbinal_data$diet_2cat[c(which(turbinal_data$diet == "Nectarivore" | turbinal_data$diet =="Frugivore"))] = "Plantivore"
turbinal_data$diet_2cat <- as.factor(turbinal_data$diet_2cat)
turbinal_data$MT_comp <- log(turbinal_data$MT.MTCH+1)
turbinal_data$ET_comp <- log(turbinal_data$ET.ETCH+1)

#-------------------------------------------------------------------------------
# Likelihood profile analysis
#-------------------------------------------------------------------------------

lambda_grid <- seq(0, 1, by = 0.05)

models <- list(
  MT_comp = MT_comp ~ wc2.1_10m_bio_7 + wc2.1_10m_bio_17 + echolocation + log(cranial_length),
  ET_comp = ET_comp ~ diet_2cat + echolocation + log(cranial_length),
  MT_SA   = log(MT + 1) ~ wc2.1_10m_bio_7 + wc2.1_10m_bio_17 + echolocation + log(cranial_length),
  ET_SA   = log(ET + 1) ~ diet_2cat + echolocation + log(cranial_length)
)

# Model fitting function

fit_gls_lambda <- function(tree, data, formula, lambda, tree_id, model_name) {

  fit <- tryCatch(
    gls(
      formula,
      correlation = corPagel(
        value = lambda,
        phy = tree,
        form = ~ species,
        fixed = TRUE
      ),
      data = data,
      method = "ML"
    ),
    error = function(e) NULL
  )

  if (is.null(fit)) {
    return(data.frame(
      model = model_name,
      tree = tree_id,
      lambda = lambda,
      logLik = NA,
      AIC = NA,
      converged = FALSE
    ))
  }

  data.frame(
    model = model_name,
    tree = tree_id,
    lambda = lambda,
    logLik = as.numeric(logLik(fit)),
    AIC = AIC(fit),
    converged = TRUE
  )
}

# -----------------------------
# Run likelihood profiles
# across all models and trees
# -----------------------------

profile_results <- bind_rows(
  lapply(names(models), function(model_name) {

    model_formula <- models[[model_name]]

    bind_rows(
      lapply(seq_along(batstrees), function(i) {

        bind_rows(
          lapply(lambda_grid, function(lam) {

            fit_gls_lambda(
              tree = batstrees[[i]],
              data = turbinal_data,
              formula = model_formula,
              lambda = lam,
              tree_id = i,
              model_name = model_name
            )

          })
        )

      })
    )

  })
)

# -----------------------------
# Add relative likelihood values
# separately for each model and tree
# -----------------------------

profile_results <- profile_results %>%
  group_by(model, tree) %>%
  mutate(
    max_logLik = max(logLik, na.rm = TRUE),
    min_AIC = min(AIC, na.rm = TRUE),
    delta_logLik = logLik - max_logLik,
    delta_AIC = AIC - min_AIC
  ) %>%
  ungroup()

# -----------------------------
# Best lambda per model per tree
# -----------------------------

best_lambda_by_tree <- profile_results %>%
  filter(converged) %>%
  group_by(model, tree) %>%
  slice_max(logLik, n = 1, with_ties = FALSE) %>%
  ungroup()

best_lambda_by_tree

# -----------------------------
# Summary of best lambda values
# -----------------------------

best_lambda_summary <- best_lambda_by_tree %>%
  group_by(model) %>%
  summarise(
    n_trees = n(),
    median_best_lambda = median(lambda, na.rm = TRUE),
    mean_best_lambda = mean(lambda, na.rm = TRUE),
    prop_lambda_0 = mean(lambda == 0, na.rm = TRUE),
    prop_lambda_le_0.05 = mean(lambda <= 0.05, na.rm = TRUE),
    prop_lambda_1 = mean(lambda == 1, na.rm = TRUE),
    .groups = "drop"
  )

best_lambda_summary

# -----------------------------
# Median likelihood profile
# -----------------------------

summary_profiles <- profile_results %>%
  filter(converged) %>%
  group_by(model, lambda) %>%
  summarise(
    median_delta_logLik = median(delta_logLik, na.rm = TRUE),
    lower = quantile(delta_logLik, 0.025, na.rm = TRUE),
    upper = quantile(delta_logLik, 0.975, na.rm = TRUE),
    .groups = "drop"
  )

facet_labels <- c(
  "ET_comp" = "ETc complexity",
  "ET_SA" = "ETc surface area",
  "MT_comp" = "MT complexity",
  "MT_SA" = "MT surface area"
)

p_summary <- ggplot(
  summary_profiles,
  aes(x = lambda, y = median_delta_logLik)
) +
  geom_ribbon(aes(ymin = lower, ymax = upper), alpha = 0.25) +
  geom_line(linewidth = 1) +
  geom_hline(yintercept = 0, linetype = "dashed") +
  facet_wrap(
    ~ model,
    scales = "free_y",
    labeller = as_labeller(facet_labels)
  ) +
  labs(
    x = "Pagel's lambda",
    y = "Relative log-likelihood",
    title = "Median likelihood profiles across trees"
  ) +
  theme_classic() +
  theme(
    plot.title = element_text(size = 14, hjust = 0.5),
    axis.title = element_text(size = 12),
    axis.text = element_text(size = 8),
    strip.text = element_text(size = 10)
  )

p_summary

#-------------------------------------------------------------------------------
# PGLS regressions for MT (likelihood profiles NOT peaking at the boundary)
#-------------------------------------------------------------------------------

#Build PGLS model

pgls_pertree <- function(model_form, tree){
  fit <- gls(model_form, correlation = corPagel(
    value = 0, phy = tree,
    form = ~ species, fixed = FALSE),
    data = turbinal_data, method = "ML")

#ANOVA
  fit_anova <- Anova(fit,type = 2)

#Tukeys for echolocation mode
  fit_tukey <- glht(fit,
    linfct = mcp(echolocation = "Tukey")
  )

  return(list(
    model = fit,
    anova = fit_anova,
    tukey = summary(fit_tukey)
  ))
}

summarize_pgls_multitree <- function(model_form, trees){

  results <- lapply(trees, function(tree){
    pgls_pertree(model_form, tree)
  })

  anova_all <- data.frame()
  tukey_all <- data.frame()

  for(i in seq_along(results)){

    # ANOVA table for one tree
    anova_tab <- as.data.frame(results[[i]]$anova)
    anova_tab$term <- rownames(anova_tab)
    anova_tab$tree <- i

    anova_all <- rbind(anova_all, anova_tab)

    # Tukey table for one tree
    tuk <- results[[i]]$tukey

    tukey_tab <- data.frame(
      tree = i,
      comparison = names(tuk$test$coefficients),
      effect_size = tuk$test$coefficients,
      p_adj = tuk$test$pvalues
    )

    tukey_all <- rbind(tukey_all, tukey_tab)
  }

  # summarize ANOVA across trees
  anova_summary <- aggregate(
    cbind(`Pr(>Chisq)`, Chisq) ~ term,
    data = anova_all,
    FUN = median
  )

  names(anova_summary)[names(anova_summary) == "Pr(>Chisq)"] <- "median_p"
  names(anova_summary)[names(anova_summary) == "Chisq"] <- "median_chisq"

  anova_sig <- aggregate(
    `Pr(>Chisq)` ~ term,
    data = anova_all,
    FUN = function(x) mean(x < 0.05) * 100
  )

  names(anova_sig)[2] <- "percent_trees_significant"

  anova_summary <- merge(anova_summary, anova_sig, by = "term")

  # summarize Tukey across trees
  tukey_summary <- aggregate(
    cbind(effect_size, p_adj) ~ comparison,
    data = tukey_all,
    FUN = median
  )

  names(tukey_summary)[names(tukey_summary) == "effect_size"] <- "median_effect_size"
  names(tukey_summary)[names(tukey_summary) == "p_adj"] <- "median_p_adj"

  tukey_sig <- aggregate(
    p_adj ~ comparison,
    data = tukey_all,
    FUN = function(x) mean(x < 0.05) * 100
  )

  names(tukey_sig)[2] <- "percent_trees_significant"

  tukey_summary <- merge(tukey_summary, tukey_sig, by = "comparison")

  list(
    anova_all = anova_all,
    tukey_all = tukey_all,
    anova_summary = anova_summary,
    tukey_summary = tukey_summary
  )
}

pgls_MTcomp_results <- summarize_pgls_multitree(models$MT_comp,batstrees)
pgls_MTSA_results <- summarize_pgls_multitree(models$MT_SA,batstrees)

#-------------------------------------------------------------------------------
# GLS regressions for ET (likelihood profiles peaking at the boundary)
#-------------------------------------------------------------------------------
pgls_zerolambda <- function(model_form, tree){

  fit <- gls(
    model_form,
    correlation = corPagel(
      value = 0,
      phy = tree,
      form = ~ species,
      fixed = TRUE
    ),
    data = turbinal_data,
    method = "ML"
  )

  # ANOVA
  fit_anova <- Anova(fit, type = 2)

  # Category means + CIs
  emm_diet <- emmeans(fit, ~ diet_2cat)
  emm_echo <- emmeans(fit, ~ echolocation)

  # Tukey tests
  fit_tukey_d <- glht(
    fit,
    linfct = mcp(diet_2cat = "Tukey")
  )

  fit_tukey_e <- glht(
    fit,
    linfct = mcp(echolocation= "Tukey")
  )

  return(list(
    model = fit,
    anova = fit_anova,

    emmeans_diet = summary(emm_diet),
    emmeans_diet_ci = confint(emm_diet),

    emmeans_echolocation = summary(emm_echo),
    emmeans_echolocation_ci = confint(emm_echo),

    tukey_d = summary(fit_tukey_d),
    tukey_d_ci = confint(fit_tukey_d),

    tukey_e = summary(fit_tukey_e),
    tukey_e_ci = confint(fit_tukey_e)
  ))
}

pgls_ETcomp_results <- pgls_zerolambda(models$ET_comp,batstrees_mcc)
pgls_ETSA_results <- pgls_zerolambda(models$ET_SA,batstrees_mcc)

#-------------------------------------------------------------------------------
# Stochastic character mapping
#-------------------------------------------------------------------------------

#fitdata (complexity values ordered according to phylogeny) will be used for phenograms and model fitting
fitdata_ETcomp <- data.frame(turbinal_data$ET_comp)
rownames(fitdata_ETcomp) <- rownames(turbinal_data)
fitdata_ETcomp <-  match.phylo.data(batstrees_mcc,fitdata_ETcomp)$data
colnames(fitdata_ETcomp) <- "ET_comp"

fitdata_MTcomp <- data.frame(turbinal_data$MT_comp)
rownames(fitdata_MTcomp) <- rownames(turbinal_data)
fitdata_MTcomp <-  match.phylo.data(batstrees_mcc,fitdata_MTcomp)$data
colnames(fitdata_MTcomp) <- "MT_comp"

#defining character vectors for stochastic character mapping
echo3vector <-  turbinal_data$echolocation
names(echo3vector) <- underscorer(turbinal_data$species)

diet2vector <- turbinal_data$diet_2cat
names(diet2vector) <- underscorer(turbinal_data$species)

#simmaps for maximum clade credibility tree
echo3simmap_mcc <- make.simmap(batstrees_mcc,echo3vector,model="ER")
diet2simmap_mcc <- make.simmap(batstrees_mcc,diet2vector,model="ER")

set.seed(123); batstrees_subsample <- batstrees[sample(seq_along(batstrees), 100)]

# Function to choose best Mk model and make simmaps
make_best_simmaps <- function(tree, map_trait, nsim = 10){

  # Fit Mk models
  fit_er <- fitMk(tree, map_trait, model = "ER")
  fit_sym <- fitMk(tree, map_trait, model = "SYM")
  fit_ard <- fitMk(tree, map_trait, model = "ARD")

  # Compare AIC
  aic_table <- data.frame(
    model = c("ER", "SYM", "ARD"),
    AIC = c(AIC(fit_er), AIC(fit_sym), AIC(fit_ard))
  )

  # Pick best model
  best_model <- aic_table$model[which.min(aic_table$AIC)]

  # Make simmaps using best model
  sims <- make.simmap(
    tree,
    map_trait,
    model = best_model,
    nsim = nsim
  )

  return(list(
    simmaps = sims,
    best_model = best_model,
    aic_table = aic_table
  ))
}

# Echolocation 3-state simmaps
echo3_results <- lapply(
  batstrees_subsample,
  make_best_simmaps,
  map_trait = echo3vector,
  nsim = 10
)

echo3simmaps <- unlist(
  lapply(echo3_results, function(x) x$simmaps),
  recursive = FALSE
)

echo3_models <- data.frame(
  tree = seq_along(echo3_results),
  best_model = sapply(echo3_results, function(x) x$best_model)
)

# Diet 2-state simmaps
diet2_results <- lapply(
  batstrees_subsample,
  make_best_simmaps,
  map_trait = diet2vector,
  nsim = 10
)

diet2simmaps <- unlist(
  lapply(diet2_results, function(x) x$simmaps),
  recursive = FALSE
)

diet2_models <- data.frame(
  tree = seq_along(diet2_results),
  best_model = sapply(diet2_results, function(x) x$best_model)
)

# Check selected models
table(echo3_models$best_model)
table(diet2_models$best_model)

#-------------------------------------------------------------------------------
# Model fitting
#-------------------------------------------------------------------------------

# MT complexity only
# Candidate set: BM1, OU1, OUM_echo

modelfit <- function(simmap1, data){

  data <- match.phylo.data(simmap1, data)$data

  BM1 <- mvBM(simmap1, data, model = "BM1",
              param = list(smean = TRUE))

  OU1 <- mvOU(simmap1, data, model = "OU1",
              param = list(root = FALSE))

  OUM <- mvOU(simmap1, data, model = "OUM",
              param = list(root = FALSE))

  AIC_values <- c(
    BM1 = as.numeric(BM1$AICc),
    OU1 = as.numeric(OU1$AICc),
    OUM_echo = as.numeric(OUM$AICc)
  )

  results <- aicw(AIC_values)
  results$AICc <- AIC_values
  results$models <- names(AIC_values)

  params <- list(
    BM1 = c(BM1$theta, BM1$sigma),
    OU1 = c(OU1$theta, OU1$sigma, OU1$alpha),
    OUM_echo = c(OUM$theta, OUM$sigma, OUM$alpha)
  )

  return(list(results, params))
}

# ET complexity only
# Candidate set: BM1, OU1, OUM_echo, OUM_diet

modelfit_joint_ET <- function(echo_simmap, diet_simmap, data){

  data_echo <- match.phylo.data(echo_simmap, data)$data
  data_diet <- match.phylo.data(diet_simmap, data)$data

  BM1 <- mvBM(echo_simmap, data_echo, model = "BM1",
              param = list(smean = TRUE))

  OU1 <- mvOU(echo_simmap, data_echo, model = "OU1",
              param = list(root = FALSE))

  OUM_echo <- mvOU(echo_simmap, data_echo, model = "OUM",
                   param = list(root = FALSE))

  OUM_diet <- mvOU(diet_simmap, data_diet, model = "OUM",
                   param = list(root = FALSE))

  AIC_values <- c(
    BM1 = as.numeric(BM1$AICc),
    OU1 = as.numeric(OU1$AICc),
    OUM_echo = as.numeric(OUM_echo$AICc),
    OUM_diet = as.numeric(OUM_diet$AICc)
  )

  results <- aicw(AIC_values)
  results$AICc <- AIC_values
  results$models <- names(AIC_values)

  params <- list(
    BM1 = c(BM1$theta, BM1$sigma),
    OU1 = c(OU1$theta, OU1$sigma, OU1$alpha),
    OUM_echo = c(OUM_echo$theta, OUM_echo$sigma, OUM_echo$alpha),
    OUM_diet = c(OUM_diet$theta, OUM_diet$sigma, OUM_diet$alpha)
  )

  return(list(results, params))
}

#-------------------------------------------------------------------------------
# Run model fits
#-------------------------------------------------------------------------------

modelfits_echo_MTcomp <- lapply(
  echo3simmaps,
  modelfit,
  data = fitdata_MTcomp
)

modelfits_ETcomp_joint <- mapply(
  modelfit_joint_ET,
  echo_simmap = echo3simmaps,
  diet_simmap = diet2simmaps,
  MoreArgs = list(data = fitdata_ETcomp),
  SIMPLIFY = FALSE
)

#-------------------------------------------------------------------------------
# Median AICc scores and weights
#-------------------------------------------------------------------------------

median_modelfit <- function(modelfits){

  model_names <- modelfits[[1]][[1]]$models

  AIC_median <- apply(
    sapply(modelfits, function(x) x[[1]]$AICc),
    1,
    median
  )

  diff_median <- apply(
    sapply(modelfits, function(x) x[[1]]$delta),
    1,
    median
  )

  AICw_median <- apply(
    sapply(modelfits, function(x) x[[1]]$w),
    1,
    median
  )

  data.frame(
    model = model_names,
    median_AICc = AIC_median,
    median_delta_AICc = diff_median,
    median_AIC_weight = AICw_median,
    row.names = NULL
  )
}

modelfits_echo_MTcomp_median <- median_modelfit(modelfits_echo_MTcomp)
modelfits_ETcomp_joint_median <- median_modelfit(modelfits_ETcomp_joint)

#-------------------------------------------------------------------------------
# Kruskal-Wallis test for AICc differences among models
#-------------------------------------------------------------------------------

test_medians <- function(modelfits){

  AIClist <- sapply(modelfits, function(x) x[[1]]$AICc)

  df <- as.data.frame(AIClist)
  df$model <- modelfits[[1]][[1]]$models

  tidy_df <- df %>%
    pivot_longer(
      cols = -model,
      names_to = "tree",
      values_to = "AICc"
    )

  kruskal.test(AICc ~ model, data = tidy_df)
}

modelfits_echo_MTcomp_wiltest <- test_medians(modelfits_echo_MTcomp)
modelfits_ETcomp_joint_wiltest <- test_medians(modelfits_ETcomp_joint)

#-------------------------------------------------------------------------------
# Percent support: models within delta AICc < 2
#-------------------------------------------------------------------------------

wincriterion <- function(difflist){
  which(difflist < 2)
}

wins <- function(modelfits){
  lapply(modelfits, function(x) wincriterion(x[[1]]$delta))
}

percentwins <- function(modelfits){

  winlist <- wins(modelfits)
  ntrees <- length(winlist)
  nmodels <- length(modelfits[[1]][[1]]$models)

  counts <- table(
    factor(
      unlist(winlist),
      levels = seq_len(nmodels)
    )
  )

  percent <- as.numeric(counts / ntrees * 100)

  data.frame(
    model = modelfits[[1]][[1]]$models,
    support_percent = percent
  )
}

modelfits_echo_MTcomp_percentwins <- percentwins(modelfits_echo_MTcomp)
modelfits_ETcomp_joint_percentwins <- percentwins(modelfits_ETcomp_joint)

# Simulated data with the winning model
# Existing dependencies not defined here: median_winparams(), modelfits_echo_ETcomp.
power_anal = function(modelfits,simmap,winner){
  params = median_winparams(modelfits)
  if (winner=='OUM'){paramlist = list(theta = c(params[1],params[2],params[3]),
                                      sigma = params[4], alpha = params[5])}
  else if (winner=='BM1'){paramlist = list(theta = params[1],
                                            sigma = params[2])}
  else if (winner=='OU'){paramlist = list(theta = params[1],
                                         sigma = params[2], alpha = params[3])}
  sim_data = mvSIM(simmap, nsim=500,model=winner,param = paramlist)
  sim_modelfits = lapply(as.data.frame(sim_data), function(col) {
    modelfit(simmap, as.matrix(col))
  })
  test_medians(sim_modelfits)
}

modelfits_echo_MTcomp_poweranal = power_anal(modelfits_echo_MTcomp,echo3simmap_mcc,'OUM')
modelfits_echo_ETcomp_poweranal = power_anal(modelfits_echo_ETcomp,echo3simmap_mcc,'OUM')

# -----------------------------------------------------------------------------
# Morphological disparity
#-----------------------------------------------------------------------------
# All species
var_MT_all <- var(turbinal_data$MT_comp, na.rm = TRUE)
var_ET_all <- var(turbinal_data$ET_comp, na.rm = TRUE)

# Nasal emitters
nasal <- subset(turbinal_data, echolocation == "Nasal")
oral <- subset(turbinal_data, echolocation == "Oral")
none <- subset(turbinal_data, echolocation == "None")

var_MT_nasal <- var(nasal$MT_comp, na.rm = TRUE)
var_ET_nasal <- var(nasal$ET_comp, na.rm = TRUE)
var_MT_oral <- var(oral$MT_comp, na.rm = TRUE)
var_ET_oral <- var(oral$ET_comp, na.rm = TRUE)
var_MT_none <- var(none$MT_comp, na.rm = TRUE)
var_ET_none <- var(none$ET_comp, na.rm = TRUE)

rel_var_MT_nasal <- var_MT_nasal/var_MT_all
rel_var_ET_nasal <- var_ET_nasal/var_ET_all
rel_var_MT_oral <- var_MT_oral/var_MT_all
rel_var_ET_oral <- var_ET_oral/var_ET_all
rel_var_MT_none <- var_MT_none/var_MT_all
rel_var_ET_none <- var_ET_none/var_ET_all

relative_variance <- rbind(
  Nasal = c(MT = rel_var_MT_nasal, ET = rel_var_ET_nasal),
  Oral  = c(MT = rel_var_MT_oral,  ET = rel_var_ET_oral),
  None  = c(MT = rel_var_MT_none,  ET = rel_var_ET_none)
)

print(round(relative_variance, 3))
```
