## Supplementary Code for "Functional tradeoffs promote morphological disparity in the bat nasal cavity": Turbinals_climates.R

```
library(terra)
library(geodata)
library(dplyr)

# Remove rows with missing latitude
locality_data <- locality_data[!is.na(locality_data$lat), ]

# Load Bioclim layers
bio <- geodata::worldclim_global(
  var = "bio",
  res = 10,
  path = "data"
)

# Convert coordinates to spatial points
pts <- terra::vect(
  locality_data,
  geom = c("long", "lat"),
  crs = "EPSG:4326"
)

# Extract climate values and combine with locality data
bio_vals <- terra::extract(bio, pts)
climate_data <- bind_cols(locality_data, bio_vals[, -1])

write.csv(climate_data, file = "climate_data.csv")
```
