## Supplementary Code for "Functional tradeoffs promote morphological disparity in the bat nasal cavity": Turbinals_climates_Hypsignathus.R

```
library(terra)
library(sf)
library(geodata)
library(dplyr)
library(tidyverse)

# Load IUCN range
iucn <- st_read("MAMMALS_TERRESTRIAL_ONLY.shp")

# Select species
sp <- iucn %>%
  filter(sci_name == "Hypsignathus monstrosus")


# Download WorldClim bioclim variables
bio <- worldclim_global(
  var = "bio",
  res = 10,
  path = "data"
)

# Convert to terra vector and match CRS
sp_vect <- vect(st_transform(sp, crs(bio)))

# Extract all bioclim values within range
vals <- terra::extract(
  bio,
  sp_vect,
  cells = FALSE
)

# Remove polygon ID column
vals <- vals[, -1]

# Summary statistics
climate_summary <- data.frame(
  Variable = names(vals),
  Mean = sapply(vals, mean, na.rm = TRUE),
  SD = sapply(vals, sd, na.rm = TRUE),
  Min = sapply(vals, min, na.rm = TRUE),
  Max = sapply(vals, max, na.rm = TRUE)
)

write.csv(climate_summary, file = "Hypsignathus_IUCN_climate.csv")
```
