## Supplementary Figures and Tables for "Functional tradeoffs promote morphological disparity in the bat nasal cavity"

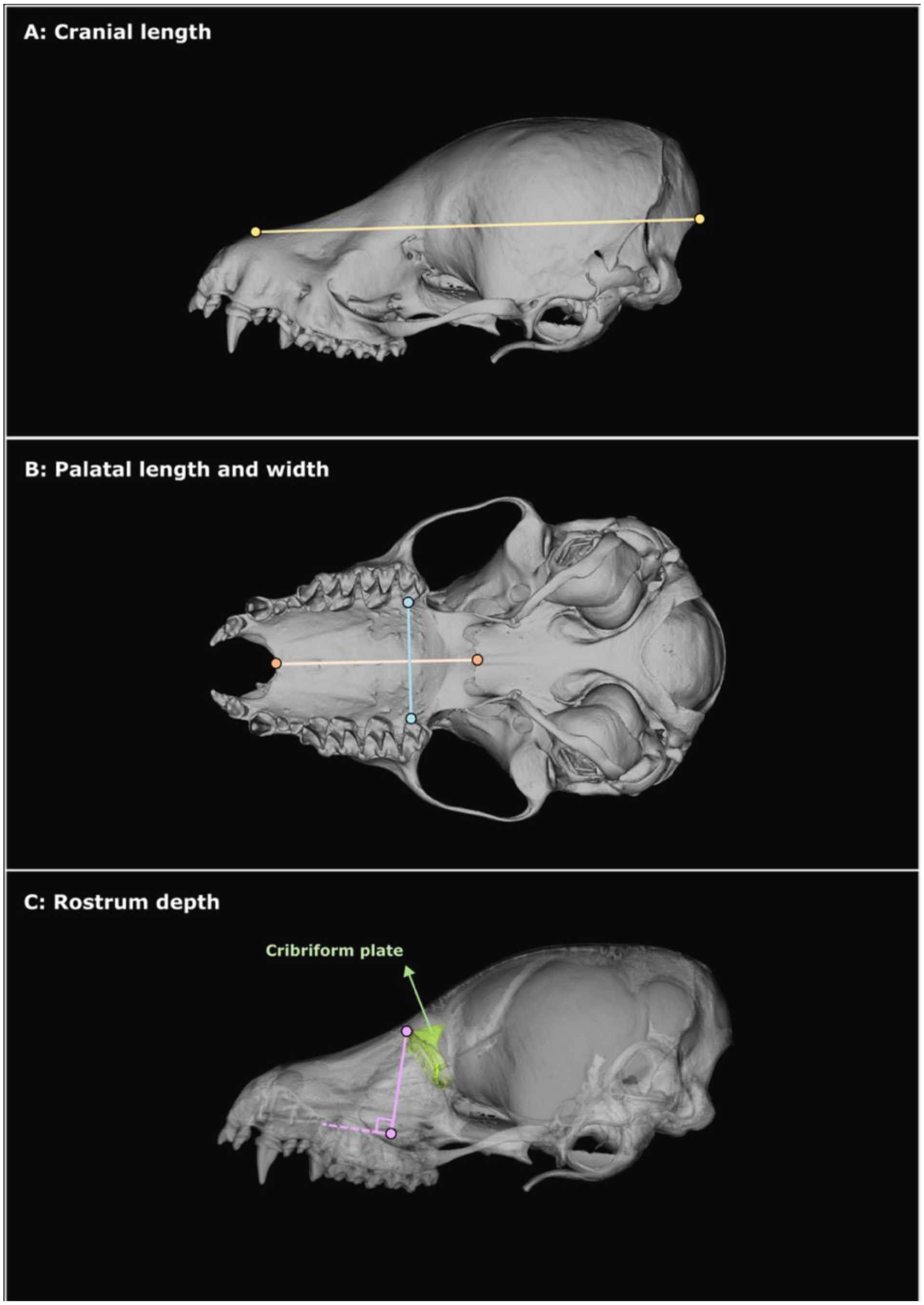

**Figure S1.** Cranial metrics used in the study. (**A**) Cranial length (distance between the anterior-most midpoint of the internasal suture (rhinion) and the dorsal-most point of the foramen magnum (opisthion). (**B)** Palatal length (distance between the anterior and posterior ends of the palatal midline) and palatal width (distance between the most lingual points of the left and right last molars in palatal view). (**C)** Rostrum depth (perpendicular distance from the most dorsal point of one half of the cribriform plate to the palatal midline)

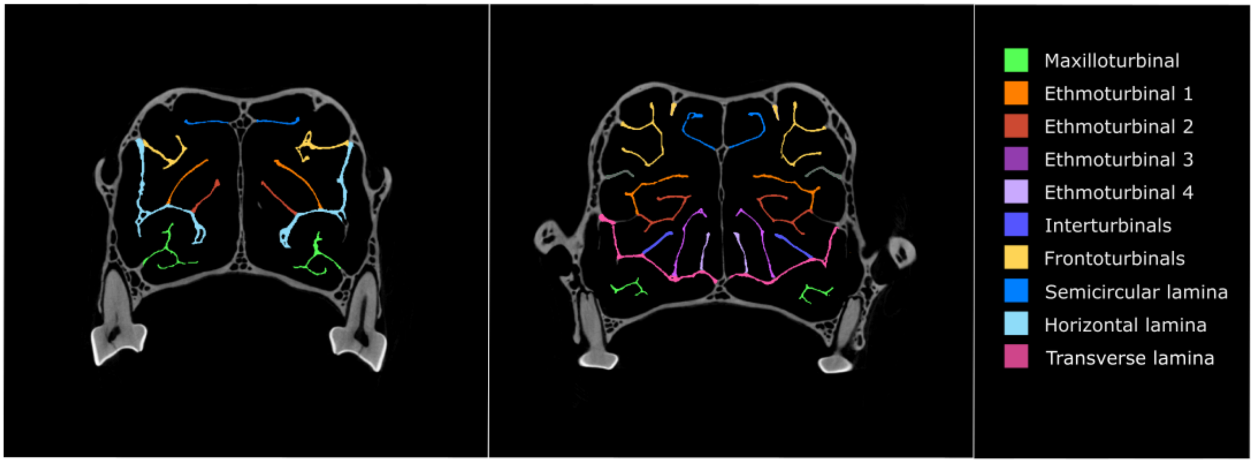

**Figure S2.** Representative coronal CT section of *Cynopterus brachyotis* showing the identities of the different turbinal elements. The ethmoturbinal complex (ETc) included the ethmoturbinals, interturbinals, frontoturbinals, and horizontal lamina, if present.

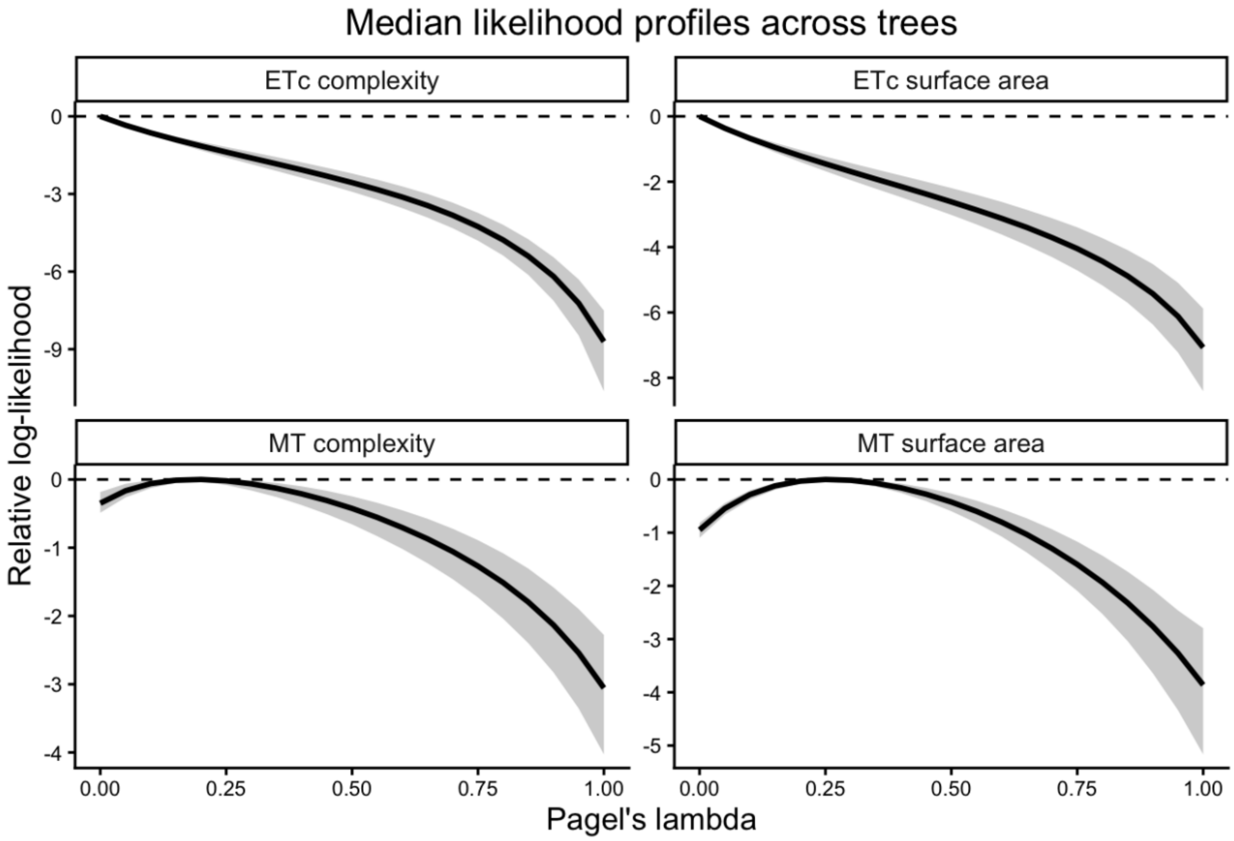

**Figure S3.** Likelihood profiles for Pagel's λ across phylogenetic trees. Profiles are shown for PGLS models of maxilloturbinal (MT) and ethmoturbinal (ETc) complexity and surface area, with log-transformed cranial length as the size covariate. For each value of λ, the solid line represents the median log-likelihood relative to the maximum log-likelihood across 1,000 phylogenetic trees obtained from VertLife (Upham et al., 2019), and the shaded ribbon represents the corresponding 2.5$^{th}$-97.5$^{th}$ percentile.

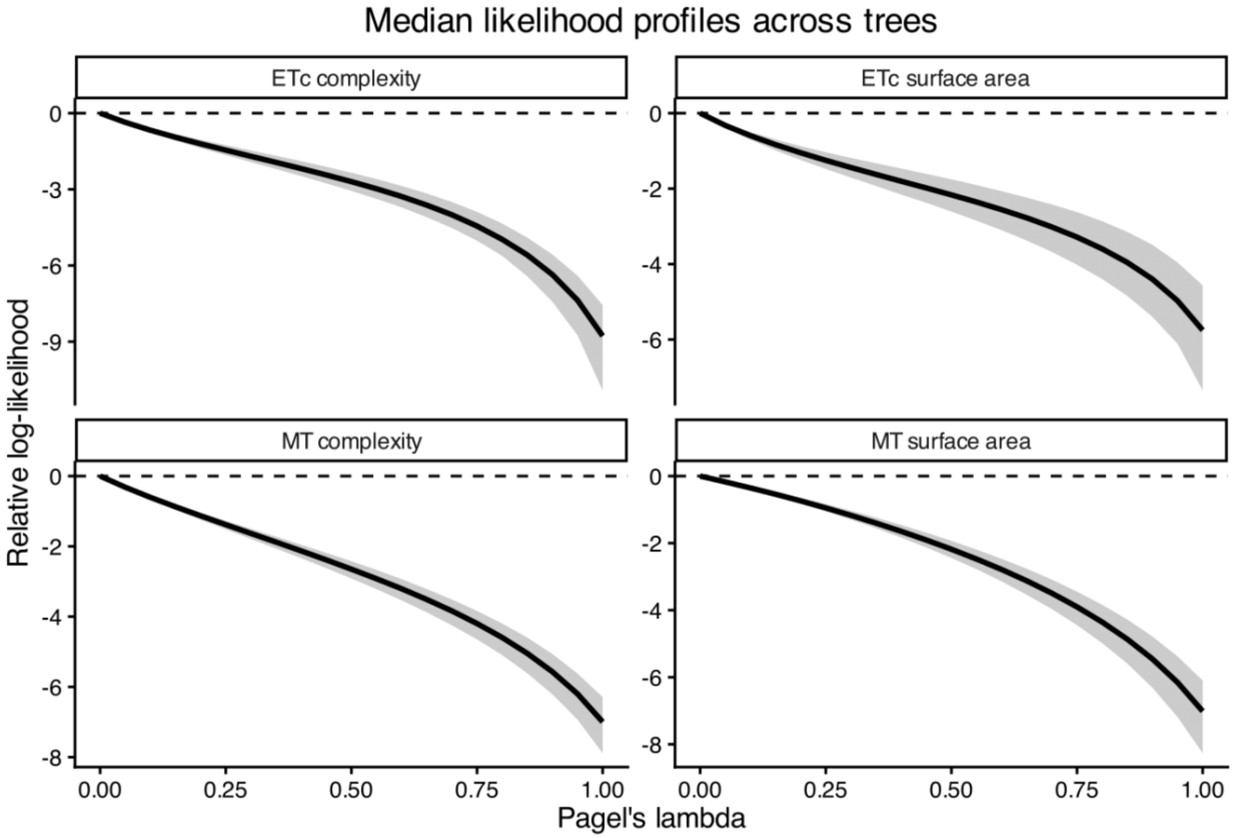

**Figure S4.** Likelihood profiles for Pagel's λ across phylogenetic trees. Profiles are shown for PGLS models of maxilloturbinal (MT) and ethmoturbinal (ETc) complexity and surface area, with log-transformed rostrum volume as the size covariate. For each value of λ, the solid line represents the median log-likelihood relative to the maximum log-likelihood across 1,000 phylogenetic trees obtained from VertLife (Upham et al., 2019), and the shaded ribbon represents the corresponding $2.5^{th}$-$97.5^{th}$ percentile.

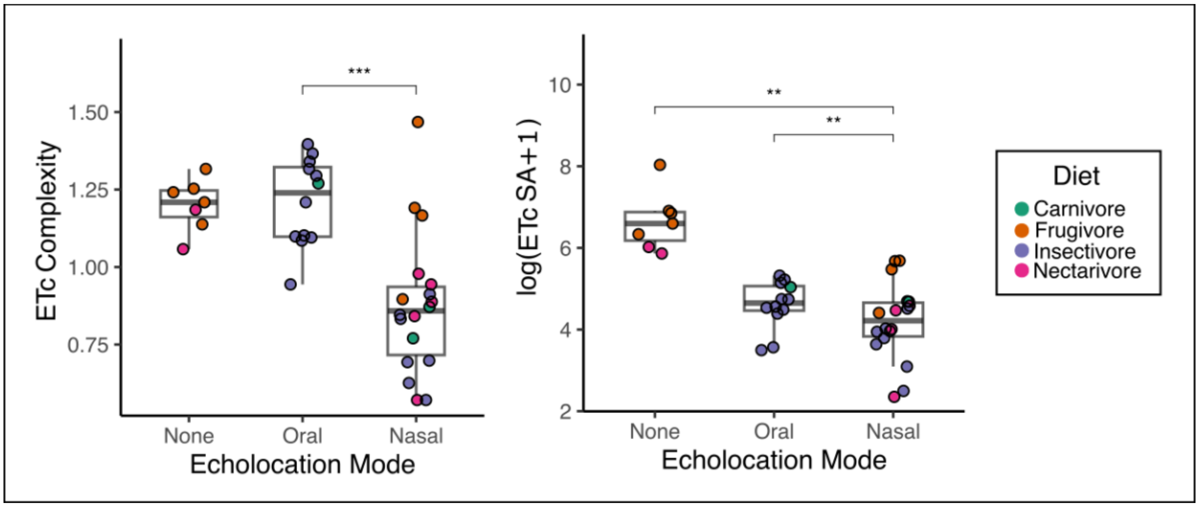

**Figure S5.** Boxplots show ethmoturbinal complex (ETc) complexity and surface area across echolocation modes, with individual species plotted as points colored according to finer-scale dietary categories. Asterisks indicate significance from ANCOVA tests of phylogenetic generalized least-squares (PGLS) models: *$p < 0.05$, **$p < 0.01$, and ***$p < 0.001$.

**Table S1.** Phylogenetic generalized least squares (PGLS) ANCOVA results for maxilloturbinal (MT) complexity and surface area using log-transformed rostrum volume as the size covariate. Models included precipitation of the driest quarter (BIO17), annual temperature range (BIO7), and echolocation mode as predictors. Pairwise comparisons among echolocation modes were conducted using Tukey's HSD post hoc test.

| **PGLS model: MT complexity ~ BIO 17 + BIO 7 + Echolocation mode + log (Rostrum volume)** | | |
|---|---|---|
| ***ANCOVA summary*** | | |
| **Predictor** | **$\chi^2$** | **p-value** |
| BIO 17 (Precipitation of the driest quarter) | 0.317 | 0.574 |
| BIO 7 (Annual temperature range) | 0.361 | 0.548 |
| log (Rostrum volume) | 23.376 | <0.001 |
| Echolocation mode | 64.987 | <0.001 |
| ***Tukey's HSD summary (Predictor: Echolocation mode)*** | | |
| **Comparison** | **Contrast estimate** | **Adjusted p-value** |
| Nasal - None | -0.485 | <0.001 |
| Nasal - Oral | -0.477 | <0.001 |
| Oral - None | -0.009 | 0.997 |
| **PGLS model: log (MT surface area+1) ~ BIO 17 + BIO 7 + Echolocation mode + log (Rostrum volume)** | | |
| ***ANCOVA summary*** | | |
| **Predictor** | **$\chi^2$** | **p-value** |
| BIO 17 (Precipitation of the driest quarter) | 0.222 | 0.638 |
| BIO 7 (Annual temperature range) | 0.048 | 0.827 |
| log (Rostrum volume) | 27.161 | <0.001 |
| Echolocation mode | 47.772 | <0.001 |
| ***Tukey's HSD summary (Predictor: Echolocation mode)*** | | |
| **Comparison** | **Contrast estimate** | **Adjusted p-value** |
| Nasal - None | -2.149 | 0.002 |
| Nasal - Oral | -2.220 | <0.001 |
| Oral - None | 0.071 | 0.994 |

**Table S2.** Phylogenetic generalized least squares (PGLS) ANCOVA results for ethmoturbinal complex (ETc) complexity and surface area using log-transformed rostrum volume as the size covariate. Models included diet and echolocation mode as predictors. Pairwise comparisons between dietary categories and among echolocation modes were conducted using Tukey's HSD post hoc test.

| | | | | |
|---|---|---|---|---|
| **PGLS model: ETc complexity ~ Diet + Echolocation mode + log (Rostrum volume)** | | | | |
| ***ANCOVA summary*** | | | | |
| **Predictor** | **$\chi^2$** | **p-value** | | |
| Diet | 6.226 | 0.013 | | |
| Echolocation mode | 38.074 | <0.001 | | |
| log (Rostrum volume) | 0.723 | 0.395 | | |
| ***Tukey's HSD summary (Predictor: Diet)*** | | | | |
| **Comparison** | **Contrast estimate** | **Lower 95% CI** | **Upper 95% CI** | **Adjusted p-value** |
| Plantivore - Faunivore | 0.209 | 0.045 | 0.374 | 0.013 |
| ***Tukey's HSD summary (Predictor: Echolocation mode)*** | | | | |
| **Comparison** | **Contrast estimate** | **Lower 95% CI** | **Upper 95% CI** | **Adjusted p-value** |
| Oral - None | 0.300 | 0.050 | 0.551 | 0.046 |
| Nasal - None | -0.146 | -0.360 | 0.069 | 0.365 |
| Nasal - Oral | -0.446 | -0.590 | -0.303 | <0.001 |
| **PGLS model: log (ETc surface area+1) ~ Diet + Echolocation mode + log (Rostrum volume)** | | | | |
| ***ANCOVA summary*** | | | | |
| **Predictor** | **$\chi^2$** | **p-value** | | |
| Diet | 1.209 | 0.272 | | |
| Echolocation mode | 11.577 | 0.003 | | |
| log (Rostrum volume) | 12.553 | <0.001 | | |
| ***Tukey's HSD summary (Predictor: Diet)*** | | | | |
| **Comparison** | **Contrast estimate** | **Lower 95% CI** | **Upper 95% CI** | **Adjusted p-value** |
| Plantivore - Faunivore | 0.359 | -0.281 | 0.998 | 0.272 |
| ***Tukey's HSD summary (Predictor: Echolocation mode)*** | | | | |
| **Comparison** | **Contrast estimate** | **Lower 95% CI** | **Upper 95% CI** | **Adjusted p-value** |
| Oral - None | -0.369 | -1.345 | 0.606 | 0.728 |
| Nasal - None | -1.067 | -1.902 | -0.232 | 0.031 |
| Nasal - Oral | -0.698 | -1.255 | -0.140 | 0.035 |
